# Single-cell RNA-seq analysis reveals aberrant CSF1 expression in disease-causing synovial fibroblasts of pigmented villonodular synovitis

**DOI:** 10.1101/2021.09.29.462128

**Authors:** Yiyong Tang, Mengjun Ma, Rujia Mi, Wenzhou Liu, Jingyi Hou, Yanjie Feng, Meijun Zhang, Menglei Yu, Fangqi Li, Yihui Song, Yixuan Lu, Yan Yan, Rui Yang

## Abstract

**Objectives:** Although the role of the CSF1/CSF1R axis in pigmented villonodular synovitis (PVNS) has been confirmed, the cells that express CSF1 and CSF1R and the underlying mechanism remain unclear. Single-cell RNA sequencing (scRNA-seq) of PVNS obtained through biopsies depicted the cellular diversity of PVNS, revealed specific CSF1/CSF1R-expressing cells and further identified novel gene expression that is associated with the development of PVNS.

**Methods:** scRNA-seq was performed on tissues obtained from the 6 biopsies of 3 patients with PVNS. Flow cytometry, immunofluorescence and western blot validated the transcriptional results, while co-culture systems revealed the cross talk between fibroblasts and macrophages.

**Results:** 8 subsets of fibroblasts and 5 subsets of macrophages were identified from the synovium of patients with PVNS and were found to be related to distinct signaling pathways. The cellular components of localized and diffuse PVNS are overall similar. Moreover, the synovium and nodule of PVNS share similar composition. The specific cells expressing CSF1/CSF1R were also identified. Other than that, unique CXCL12^+^CSF1^+^ fibroblasts were revealed to attract macrophages as disease-causing synovial fibroblasts, leading to the formation of masses in PVNS.

**Conclusions:** PVNS consists of macrophages, fibroblasts, T cells, endothelial cells and mast cells. Among them, the CSF1-expressing fibroblasts appeared to be tumor-like cells that attract macrophages, subsequently forming tumor-like mass in PVNS. This paves the path for novel treatments of PVNS by targeting CXCL12^+^CSF1^+^ fibroblasts and the CXCL12-CXCR4 axis.

## Introduction

Pigmented villonodular synovitis (PVNS), i.e. tenosynovial giant cell tumor (TGCT),^1^ is a rare, proliferative disease affecting the synovial joints or tendon sheaths in young adults aged 20 to 40 years old^2,3^. PVNS can be classified as either localized or diffuse, and usually occurs in weight-bearing joints, with the highest occurrence in the knees (66%-80%)^4^. Although arthroscopic excision remains the mainstay of treatment, diffuse PVNS is prone to relapse even after total synovectomy, with a recurrence rate of approximately 50%^5^.

As the name implies, PVNS is characterized by haemosiderin deposition, villonodular synovial hyperplasia and synovitis. There are two types of PVNS: the localized or nodular form (where the lesion involves only one area of the joint) and the diffuse form (where the entire lining of the joint is involved)^6^. Histologically, PVNS is composed of fibroblasts, macrophages, multinucleated giant cells, lymphocyte, siderophages, among others^7^. The pathogenesis of PVNS is unclear, while its categorization either as inflammatory or neoplastic disease remains controversial^8,9^. More recently, evidence of autonomous growth, malignant transformation and cytogenetic aberrations seem to lean toward a neoplastic origin of PVNS^10–12^.

West et al. proposed a landscape effect in PVNS, whereby the overexpression of colony-stimulating factor 1 (CSF1) by a small minority of neoplastic cells leads to the recruitment of CSF1 receptor (CSF1R)-expressing macrophages, resulting in the formation of tumor mass^11^. This encourages the clinical application of CSF1R inhibitors in PVNS therapy with satisfactory outcomes yielded^13–16^. CSF1 is a multifunctional protein that promotes the migration, survival and differentiation of macrophages and their precursors through CSF1R^17^. One of the CSF1R monoclonal antibody, RG7155, could effectively deplete CSF1-differentiated macrophages characterized by the expression of CSF1R and CD163, and subsequently reduced the tumor burden of PVNS patients^18^. However, the application of CSF1R inhibitor alone does not solve the problem entirely. Firstly, CSF1R inhibitor not only inhibits the proliferation of pro-tumor macrophages, but also prevents the survival of anti-tumor macrophages, which may explain the limited effect of such treatment in some PVNS patients. Secondly, the CSF1R inhibitor only interrupts the CSF1/CSF1R axis and does not stop the continuous rise of CSF1. Once discontinued, the overexpressed CSF1 will again recruit macrophages to accumulate and proliferate, leading to probable recurrence^14^. Even though West et al. also proposed chromosomal translocation involving COL6A3 and CSF1 as the cause of CSF1 overexpression^11^, such chromosomal translocation was reported to be absent from some PVNS patients^19^. To date, the specific mechanism of CSF1 overexpression in PVNS remains unclear. This missing puzzle might be the key to unlock novel PVNS treatment, and the identification of cells that highly-express CSF1 might be the prerequisite to elucidate this mechanism.

Single-cell RNA sequencing (scRNA-seq) has the advantage of effectively distinguishing intercellular heterogeneity, which makes it an ideal method to identify cells with highly-expressed CSF1 in PVNS. In the current study, markers were used to define specific CSF1-expressing fibroblast subset that recruits and promotes the polarization, migration of pro-tumor macrophages. Further analysis of the subset showed that CSF1 overexpression was accompanied by a significant increase in the expression of CXCL12, making CXCL12 a potential target to curb the effects of disease-causing synovial fibroblasts on macrophages.

## Methods

### Clinical tissue specimens and cell isolation

This study was approved by the Ethics Committee of the Sun Yat-sen Memorial Hospital, Sun Yat-sen University. During the arthroscopic surgeries, a total of 3 nodular tissues and 3 synovial tissues were obtained from 2 diffuse-type PVNS patients and 1 localized-type PVNS patient. Thereinto, the synovial tissue that collected from the localized-type PVNS patient was normal and taken as control. All patients did not receive anticancer treatments like chemotherapy or radiotherapy before tissue sampling. And the clinical characteristics of these patients were listed in figure S1B.

The tissues were clipped and cultured in medium with 0.02% collagenase type I at 70 rpm/min on a 37 □ constant temperature shaking incubators for 4-5 h until the tissues were filamentlike. After centrifugation at 1000 rpm for 5 minutes, the supernatant was removed and the cell pellets was cultured in DMEM supplemented with 10% fetal bovine serum (FBS) in a 37 °C, 5% CO_2_ incubator. After the third passage, the non-fibroblasts cells completely disappeared from these culture systems, and the remaining cells were primarily synovial fibroblasts.

### scRNA-seq library construction and sequencing

Sequencing libraries were constructed according to a modified single-cell tagged reverse transcription (STRT) protocol as our previously reported^20^. In brief, after the samples were isolated, a 25 nucleotide (nt) oligo (dT) primer anchored with an 8 nt unique barcode and an 8 nt unique molecular identifiers (UMI) were added onto the RNA ends through reverse transcription. Then, beads with relative barcodes were added to saturation for pairing with the cells in the microwells. Cell lysis buffer was added to hybridize polyadenylated RNA molecules to the beads. Then, the beads were collected into a single tube for reverse transcription. After synthesis and 18 cycles of amplification, sequencing libraries were constructed by random priming PCR to enrich the 3’ end of the transcripts that were anchored with cell label and UMI, and then submitted for 150 bp paired-end sequencing on the Illumina HiSeq 4000 platform.

### scRNA-seq data analysis

Upon constructing the sequencing libraries and completing the 150 bp paired-end sequencing, the adaptor sequence was filtered to remove low-quality reads in fastp with default parameters. Next, single cell transcriptome analysis was performed with the help of UMI-tools to check the cell barcode whitelist. The UMI-based data was mapped against the human genome (Ensemble version 91) obtained from the UMI-tools standard pipeline to determine the UMI count of each sample with STAR. Down sampling was done to minimize sample batch based on the mean read per cell of each sample and a cell expression table with a sample barcode was achieved. Cells containing over 200 expressed genes and a mitochondrial UMI rate of below 20% passed the cell quality filtering, whereas mitochondrial genes were removed. Next, normalization and regression were applied on the expression table based on the UMI count of each sample and the different sequencing library of each sample so that scaled data can be obtained with the Seurat package (version 2.3.4, https://satijalab.org/seurat/) in default settings. Next, Principal Component Analysis (PCA) was constructed to reduce dimensions of the integrated data, and PC1 to PC30 were used to identify clusters (resolution=0.01), and UMAP (uniform manifold approximation and projection) algorithm was used for visualization. FindAllMarkers function was performed based on the bimod algorithm of R package Seurat to find differentially expressed genes (DEGs) in the specified cell-types. Fold changes of ≥ 1.25 and p < 0.05 were considered as statistically significant.

Cell-cell communication analysis was performed using CellPhoneDB (version 2.1.1), the open database for ligands, receptors and their interactions, according to the manufacturer’s protocol (https://www.cellphonedb.org/). Cell-cell communications with p-value < 0.01 were considered as significant and were selected. The outcome of CellPhoneDB was applied to gene enrichment analysis with GO: BP database as the basis.

### Western blot analysis

PVNS nodular and synovial samples were harvested and lysed with RIPA buffer (1% NP-40, 0.5% sodium deoxycholate, 0.1% SDS, 10 ng/ml PMSF, 0.03% aprotinin, and 1 μM sodium orthovanadate) for 30 min on ice and the supernate was retained. The protein was separated using 12% SDS-PAGE and transferred onto PVDF membranes. The PVDF membranes were blocked with skim milk for 1 h, before incubating with primary antibodies for 12 h and secondary antibodies for 1 h respectively. Proteins were detected and qualified with chemiluminescent detection reagents and films. The primary antibodies used in this study were CSF1 (ab52864, Abcam), CSF1R (ab254357 Abcam) and GAPDH (RM2002, Ray antibody); while the goat anti-rabbit IgG (BA1054, BOSTER) and the goat anti-mouse IgG (BA1050, BOSTER) was used as the secondary antibody.

### Hematoxylin-eosin (HE) staining

After the removal of excess yellow adipose tissue, the synovial and nodular samples were scissored into 4 mm segments and fixed in 4% paraformaldehyde, then after getting through the gradient ethanol dehydration, pure xylene transparency. The sections were then embedded with paraffin and cut into 5 μm thin slices after getting through dehydration by gradient ethanol series. Next, dewaxing took place in the order of xylene, ethanol and distilled water, each for 5 min, after which HE staining was carried out for 20 min. Following that, the colors were separated with 75% hydrochloric acid alcohol for 10 s, washed in slow double-distilled water for 3 min, dipped in 95% alcohol for 1 min, and dyed with eosin for 1 min. The concentration of alcohol, dimethylbenzene, and neutral gum were set at a gradient series for dehydration, hyalinization and sealing. The sections were finally observed under a microscope (Lecia DM) at 200 × magnification.

### Immunohistochemistry (IHC) staining

Paraffin-embedded tissue sections were dewaxed, rehydrated, and treated with peroxidase and pepsin for 20 min and 30 min, respectively. Then, the sections were incubated with the primary antibodies CD68 (ab125212, Abcam), CD163 (49553, SAB), PDPN (sc-376695, Santa Cruz), FAP (BM5121, Boster), and CD34 (ab185732 Abcam) for 1 d, and secondary antibody (G1216, Servicebio) for 40 min. Next, the Histostain-Plus Kit (CW2069S, CWB) was used in accordance to the manufacturer’s instructions. The sections were finally observed under a microscope (Lecia DMi8) at 200 × magnification. Dense, dark particles represent the expression and the location of proteins CD68, CD163, PDPN, FAP and CD34.

### Immunofluorescence (IF) staining

Nodular and synovial cryostat sections of PVNS were fixed and cut into 4 μm thin slices. Then, the target gene was stained with the following antibodies: CSF1-mRNA (axl-FISHcsf1, axl-bio), CSF1R (ab254357, Abcam), PDPN (376695, Santa Cruz), CD163 (ab156769, Abcam); and the secondary antibody anti-mouse IgG (H+L), F(ab’)2 Fragment (Alexa Fluor® 488 Conjugate) (4408, CST), anti-mouse IgG (H+L), F(ab’)2 Fragment (Alexa Fluor® 555 Conjugate) (4409, CST), anti-rabbit IgG (H+L), F(ab’)2 Fragment (Alexa Fluor® 488 Conjugate) (4412, CST), Anti-rabbit IgG (H+L), F(ab’)2 Fragment (Alexa Fluor® 555 Conjugate) (4413, CST) were used. Nuclei were subsequently visualized by DAPI staining (0215757410, MP). The sections were finally observed under the microscope camera system (Olympus IM, USA) and the intensities of immunofluorescence were quantitated by the ImageJ software.

### Macrophage isolation and differentiation

Macrophages from peripheral blood mononuclear cell (PBMC) have the potential to differentiate into M1 and M2 macrophages. In this study, PBMC were isolated from peripheral blood with the classical monocyte isolation kit (LTS1077-1, TBD, Tianjin Haoyang). Then the isolated monocytes were differentiated into macrophages (M0 macrophages) after stimulation with 20 nM CSF-1 for 7 d. Subsequently, the M0 macrophages were co-cultured with fibroblasts that were separated from the normal or PVNS synovial samples. These fibroblasts were seeded in the upper insert of a six-well Transwell plate (0.4 μm pore size, Corning), while the M0 macrophages were seeded in the lower chamber. After 48 h, the macrophages were harvested for further identification with flow cytometry assay.

### Flow cytometry assay

The tissue or cell suspension was incubated with CD68-FITC (333806, Biolegend), CD163-PE (556018, BD), CD163-APC (333609, Biolegend), PDPN-FITC (ab205333, Abcam), and CD86-PE (305406, Biolegend) before proceeded for identification by the advanced analytical flow cytometer (BD FACSVerse). A sample was prepared without antibody incubation as the negative control. The ratio of M1 macrophage population was identified by CD86, M2 macrophages by CD 163 and fibroblasts by PDPN.

### Statistical analysis

Data were analyzed using SPSS Statistics 26 (SPSS, Chicago, IL). All data are expressed as the mean ± SD. Student’s *t* test and one□ way ANOVA were used to determine the differences between 2 groups. *P* < 0.05 is considered as statistically significant.

## Result

### Single-cell profiling of human PVNS lesions and normal synovial tissues

To determine the cell subsets and gene expression profile of PVNS lesions and normal synovial tissues, 6 tissues of 3 PVNS patients were collected according to the modified STRT strategy to perform scRNA-seq (figure 1A). The lesions of patients #1 and #2 were located in the knee joint, while the lesions of patient #3 was located in the ankle joint (figure 1B). Their nodular and synovial samples were respectively labeled as N1, N2, N3 and S1, S2, S3. Patient #1 and #3 were diffuse-type PVNS, while patient #2 was localized-type PVNS with only one nodule in the joint cavity, the others were normal tissues. Therefore, the sample S2 was the normal synovium that served as the control in this research (figure S1B). The nodular and synovial samples were divided into five cell clusters based on unbiased clustering: macrophages, fibroblasts, T cells, endothelial cells and mast cell (figure 1C, S2B). There was no preference in terms of sample distribution in each cluster (figure 1C, S1C). Heatmap showed differential gene expression among the five cell clusters (figure 1D). The marker genes of each cluster were: CD14, CD68, CD86, CD163 and CYBB for macrophages; FAP, PDPN, THY1 and COL1A1 for fibroblasts; CD3E, CD3D and NKG7 for T cells; PECAM1, CD34 and VWF for endothelial cells; and TPSAB1, CPA3 and MS4A2 for mast cells (figure 1E, S2A).

**Figure 1.**
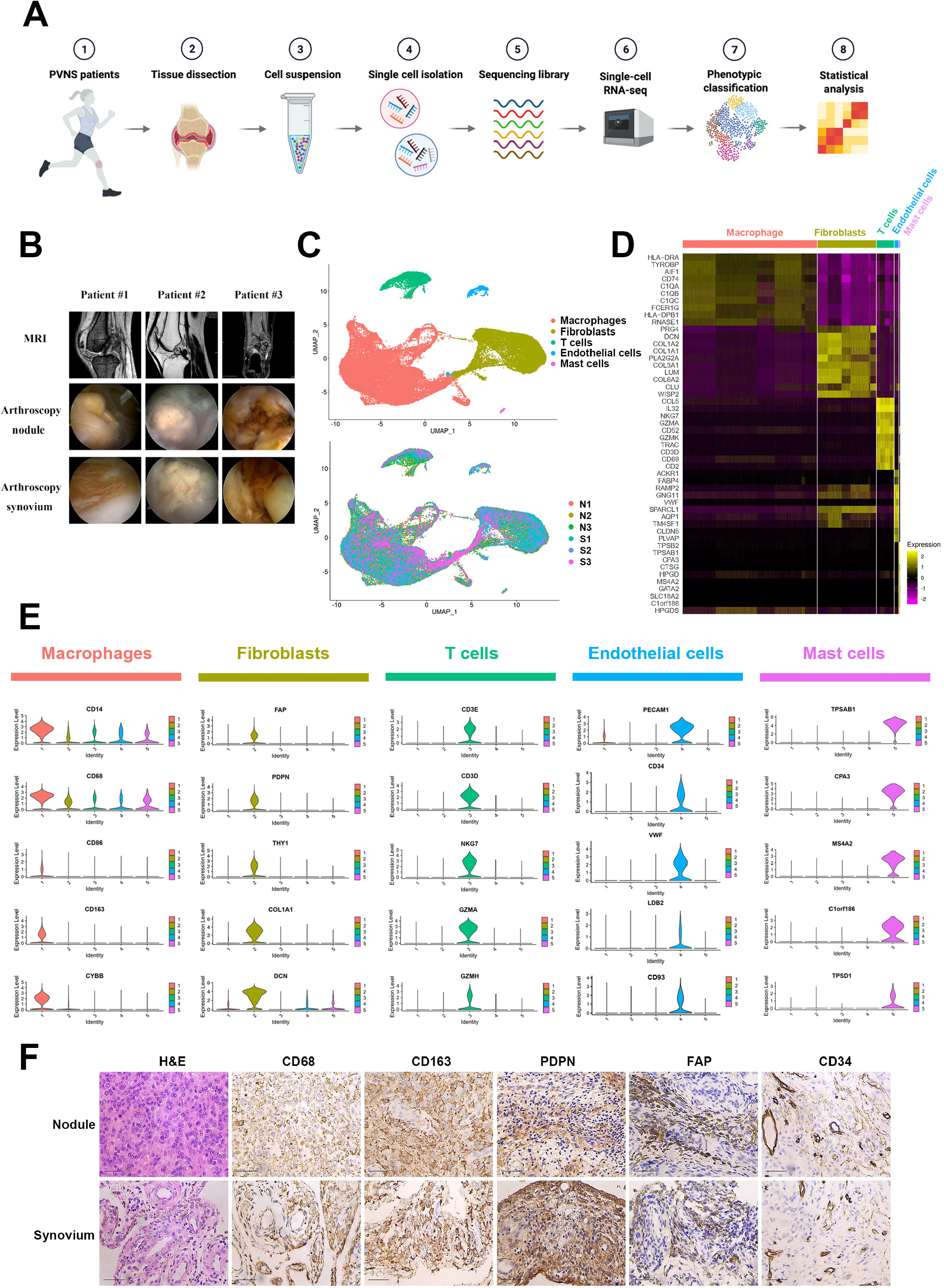
Single-cell profiling of PVNS lesions and normal synovial tissues. (A) Schematic workflow of the experimental strategy. (B) MRI and arthroscopy images of the three PVNS patients that involved in this study. (C) Profiles of the 62,898 cells that were extracted from the nodule (41,252 cells) and the synovium (21,646 cells), and are shown visually as UMAP plots. Each cell is color-coded according to its sample origin (left panel) and associated cell type (right panel). (D) Heatmap reveals the scaled expression of DEGs for each cluster. (E) Violin plots have shown the gene expression of representative candidate marker genes for the different cell types in UMAP plots. (F) H&E and IHC staining of CD68, CD163, PDPN, FAP and CD34 in the nodular and synovial samples. Dark particles represent hybrid sites.

In order to further clarify the distribution of each cell cluster in the nodular and synovial samples, IHC staining were used to identify the expression level of marker genes: CD68 and CD163 for macrophages; PDPN and FAP for fibroblasts; and CD34 for endothelial cells (figure 1F). Despite the differences in morphological character between nodular and synovial lesion, there was no preference in the distribution of macrophages, fibroblasts and endothelial clusters (figure S1C). These results indicate that the nodules and synovium of PVNS have similar composition and even the same cellular origin.

### Synovial fibroblasts were the main source of CSF1 in PVNS

To elucidate the location and expression of CSF1 and CSF1R, the expression patterns of all five clusters were analyzed. CSF1 was mainly found in the fibroblasts and mast cells, whereas CSF1R was mainly located in the macrophages (figure 2A). These results suggested that possible communication bridges between fibroblasts, mast cells and macrophages.

**Figure 2.**
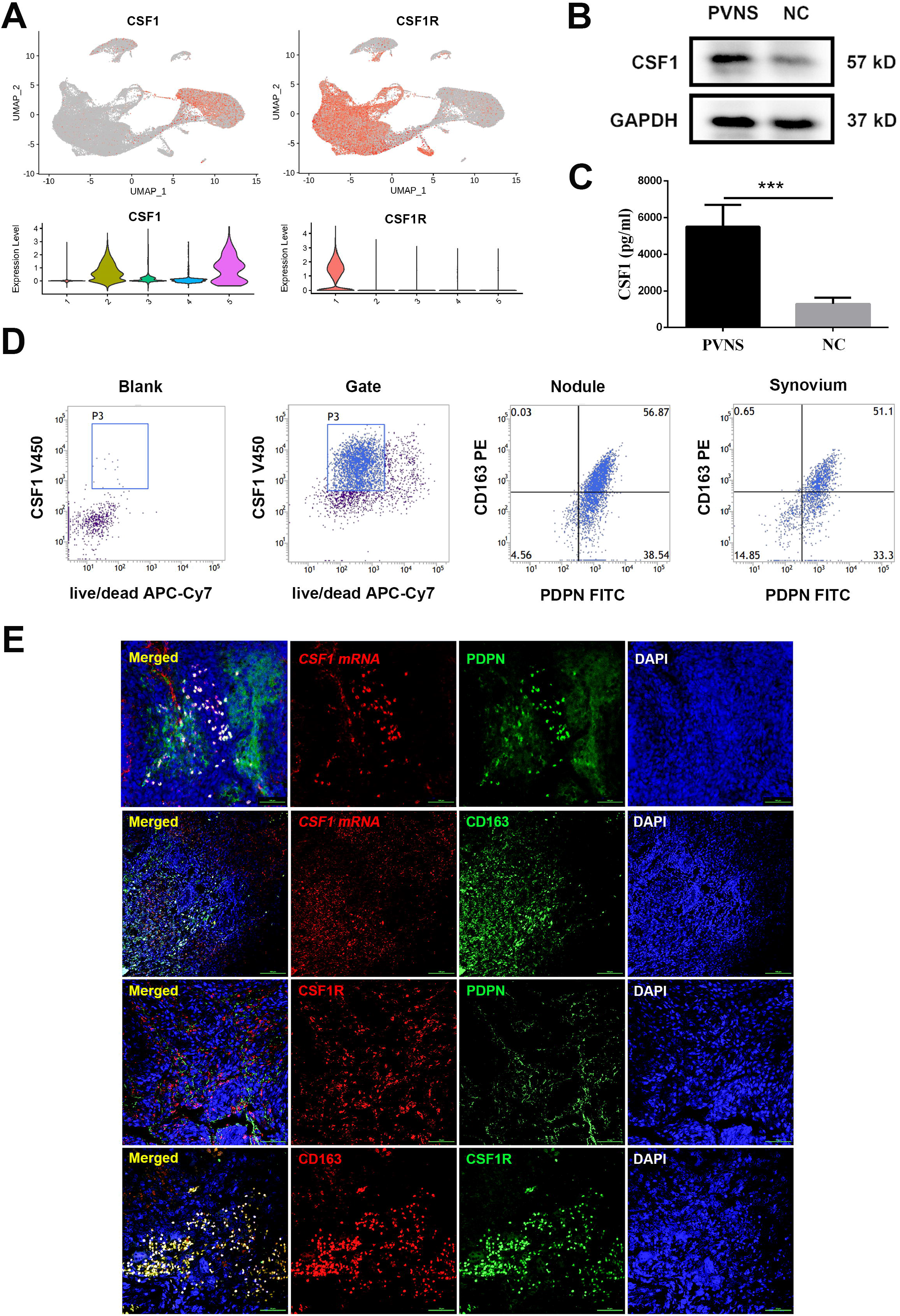
Identification of the location and expression of CSF1 and CSF1R in PVNS. (A) The location of CSF1 and CSF1R in the UMAP plot. Violin plot has shown the CSF1 mostly expressed in fibroblasts and mast cells, CSF1R mainly expressed in macrophages. (B) (C) The results of western blot showed that compared to the normal synovial tissue, CSF1 were higher expressed in PVNS lesions. All data are expressed as the mean ± SD. ***, P < 0.001. (D) The results of flow cytometry showed that the different proportion of CSF1-expressing cells in the nodular and synovial samples of PVNS. CD163 is a marker gene of macrophage, whereas PDPN is a marker gene of fibroblasts. (E) The results of IF staining showed that the CSF1 was mostly expressed in PDPN^+^ fibroblasts but not in CD163^+^ macrophages, whereas CSF1R was expressed in CD163^+^ macrophages but not in PDPN^+^ fibroblasts.

Therefore, we isolated and analyzed the nodule and synovial samples *in vitro* to identify the specific cell expression of CSF1 and CSF1R. Compared with the normal synovium tissue, a higher level of CSF1 was found in the PVNS lesion (figure 2B, 2C). Furthermore, flow cytometry showed that fibroblasts accounted for the majority of CSF1-expressing cells both in nodule and synovium of PVNS (figure 2D). Meanwhile, IF staining results revealed that CSF1 was mostly expressed in PDPN^+^ fibroblasts but not in CD163^+^ macrophages, whereas CSF1R was expressed in CD163^+^ macrophages but not in PDPN^+^ fibroblasts (figure 2E). Taken together, these results indicated that disease-causing CSF1^+^ fibroblasts mainly express and secrete CSF1, which may recruit macrophages and promote their proliferation and differentiation, thereby forming tumor-like mass in PVNS.

### CXCL12^+^CSF1^+^ fibroblasts were the disease-causing synovial fibroblasts in PVNS

The previous results suggested that fibroblasts were the main source of CSF1 in PVNS, so we further divided fibroblast subclusters in order to identify the disease-causing fibroblasts. The fibroblasts were divided to 8 subpopulations using unbiased clustering (figure 3A). Noticeably, subclusters 1, 2, 3, 8 mainly expressed CSF1 which may act as the disease-causing synovial fibroblasts in PVNS (figure 3D) and the number of subcluster 1 was the largest (figure 3A). Compared to the normal synovium (sample S2), PVNS lesions contained a unique subcluster 8 (Figure 3B, S2D, S2E). Furthermore, in addition to the well-known fibroblast marker genes of FAP, PDPN and THY1, subclusters 1, 2, 3 and 8 also specifically overexpressed TNFAIP6 and CTGF (figure 3C). Therefore, it was speculated that these 4 subclusters were made up of fibroblasts derived from the synovial sub-lining^21^. Besides the commonly overexpressed CSF1 gene, some other genes were also observed to be differentially expressed in these 4 subclusters with significance: ANGPTL1 and CXCL12 in subcluster 1; DEFB1 and TSG6 in subcluster 2; AIF1 and C1QB in subcluster 3; CST7, NKG7 and a small amount of CSF1R in subcluster 8 (figure 3D). Then the subclusters 1, 2, 3, 8 could be separately defined as CXCL12^+^CSF1^+^ fibroblasts, TSG6^+^CSF1^+^ fibroblasts, AIF1^+^CSF1^+^ fibroblasts, CST7^+^CSF1^+^ fibroblasts. And the subclusters 4, 5, 6, 7 could be together defined as CSF1^-^ fibroblasts. The heatmap showed that the expression of the marker genes of each subcluster in each subpopulation (figure 4E). Enrichment analysis was carried out on the marker genes of each subcluster which imply the different effects of each cell subpopulation (figure 4F). Specifically, subcluster 1 was associated with extracellular matrix organization and cartilage development; subcluster 2 was enriched in oxidative stress; subcluster 3 was related to inflammatory response; while subcluster 8 was associated with lymphocyte activation and differentiation. In addition, since subcluster 1 has the highest proportion, taken together, these data indicated that the subcluster 1, CXCL12^+^CSF1^+^ fibroblasts may dominate the pathogenesis and progression of PVNS.

**Figure 3.**
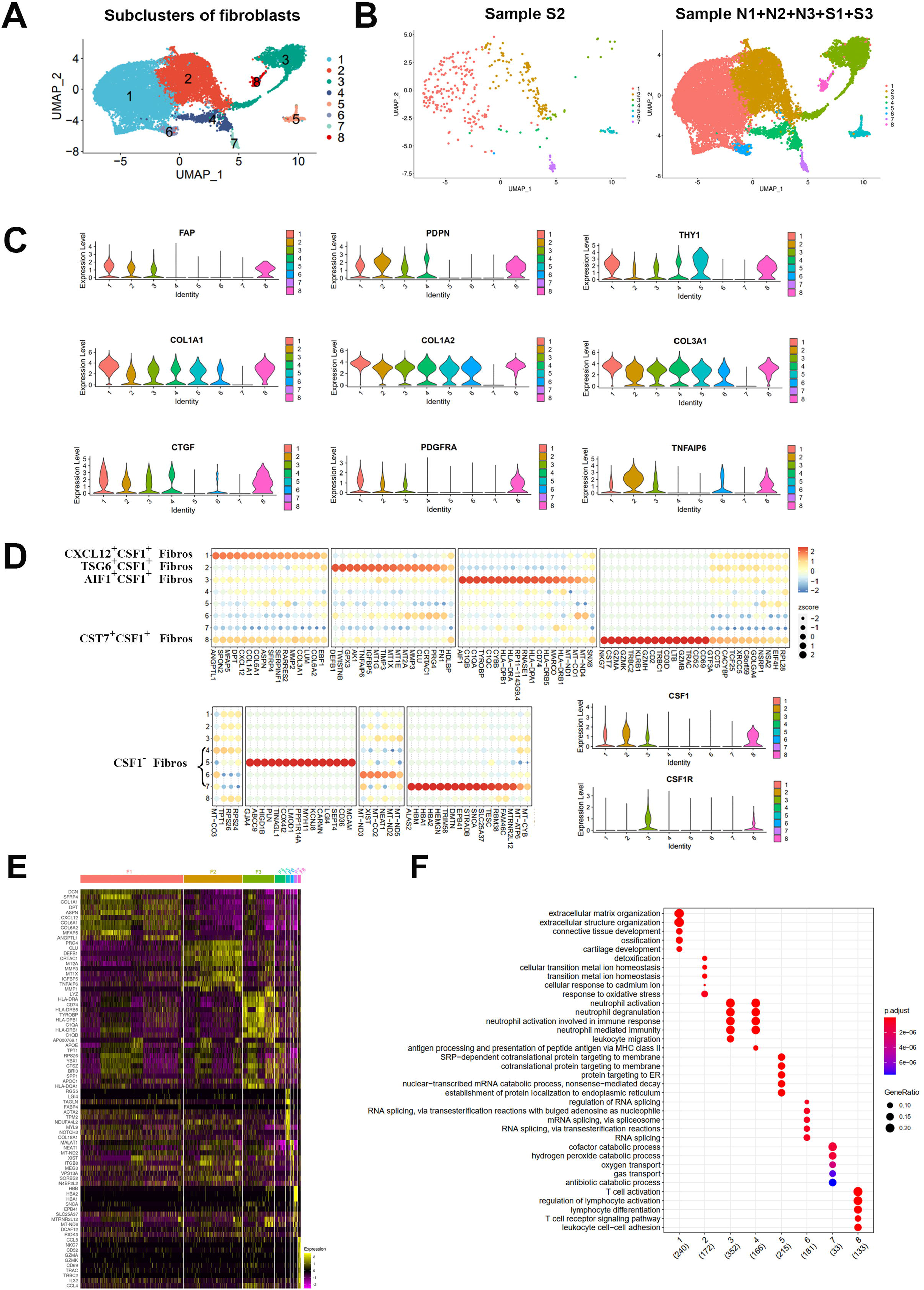
Subcluster CXCL12^+^CSF1^+^ fibroblasts were the disease-causing synovial fibroblasts in PVNS. (A) The fibroblasts were divided to 8 subclusters and visually represented as UMAP plots. (B) The proportion of fibroblast subclusters respectively in normal synovial tissue (Sample S2) and PVNS lesion (Sample N1+N2+N3+S1+S3). (C) Violin plots showed the gene expression of representative marker genes in the fibroblast subclusters. (D) The top DEGs in the fibroblast subsets. Violin plots represented the expression of CSF1 and CSF1R in the different subclusters. Subclusters 1, 2, 3, 8 mainly expressed CSF1 and subclusters 3, 8 specifically expressed CSF1R. (E) The heatmap showed that the expression of the marker genes of each subcluster in each subpopulation. (F) Enrichment analysis was carried out on the marker genes of each subcluster which imply the different effects of each cell subpopulation. The number of marker genes was indicated in brackets.

**Figure 4.**
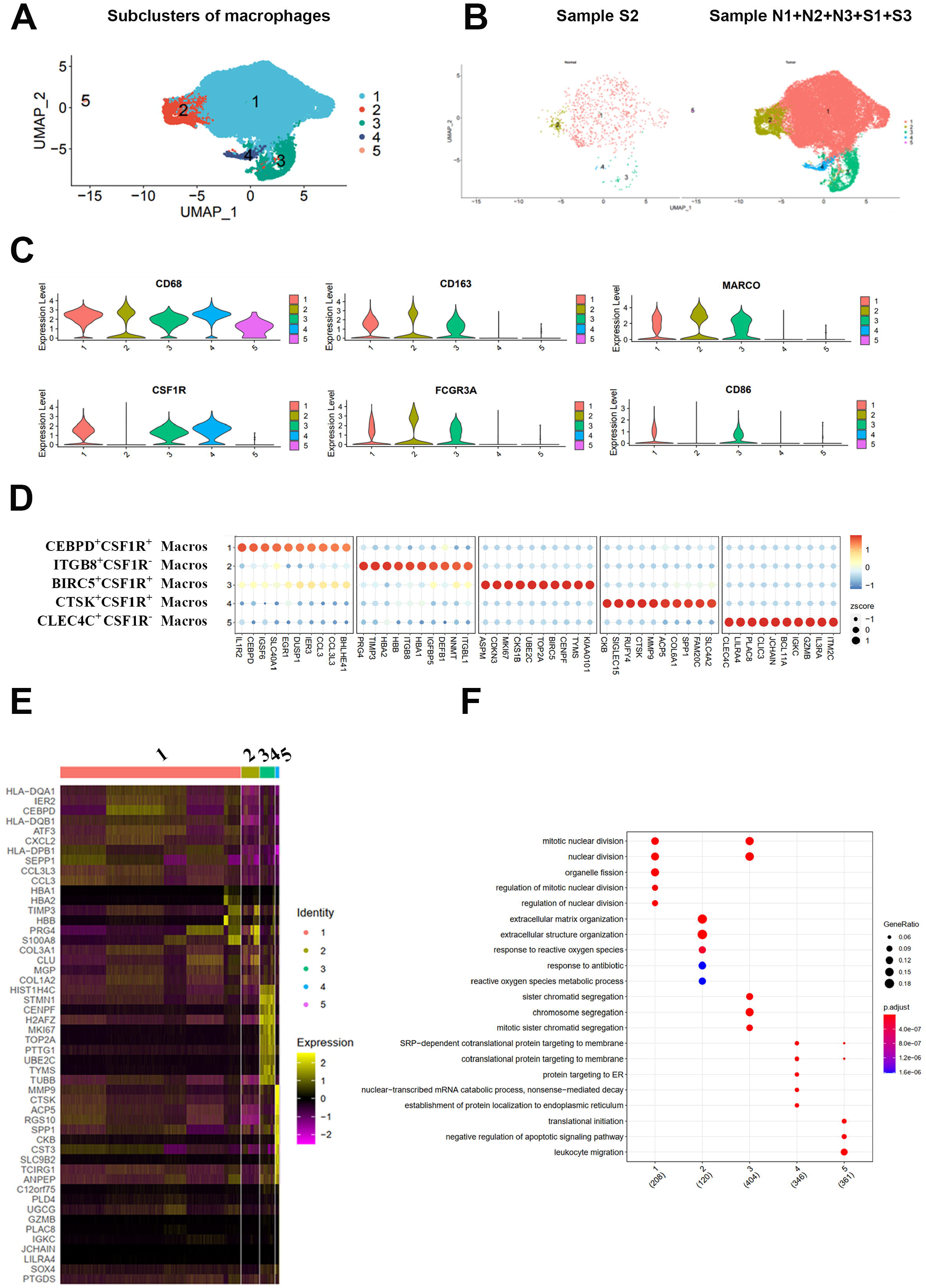
CD163^+^CSF1R^+^ macrophages may interact with CSF1^+^ fibroblasts via the CSF1-CSF1R axis in PVNS. (A) Profiles of five subclusters of macrophages which visually represented as the UMAP plots. (B) The proportion of five subclusters of macrophages in PVNS lesions and normal synovial tissues. There was no significant different distribution between them. (C) Violin plots showed that the expression of representative marker genes in the five subclusters. (D) The top ten DEGs in the macrophage subclusters. (E) The heatmap showed the detailed patterns of DEGs in five subclusters. (F) GO enrichment analysis was performed on the DEGs in five subclusters to verify their possible functions. The number of marker genes was indicated in brackets.

### CD163^+^CSF1R^+^ macrophages maybe the effector cells of CSF1^+^ fibroblasts during the PVNS tumor formation

According to the results of first UMAP plots and *in vitro* assays, CSF1R was found to be mainly expressed in macrophages. Therefore, we further divided the macrophage and tried to clarify the function of CSF1R^+^ macrophages. Unbiased clustering divided the macrophages into 5 subclusters (figure 4A) and there was no significant different distribution between PVNS lesions and normal synovial tissue (figure 4B, S2C). The number of subcluster 1 was the largest (figure 4A). In addition to the common marker gene of macrophage, CD68, subclusters 1, 2, 3 expressed CD163, which was a marker gene of M2 macrophages. Hence, subclusters 1, 2, 3 represented the M2 macrophages. However, subclusters 4, 5 lacked markers for both M1 and M2, raising speculation of their possible relationship to the M0 macrophages or the other subsets similar to macrophages. Noticeably, the CSF1R was mainly found in subclusters 1, 3 and 4, which suggested that CSF1R mostly located on the M2 macrophages (figure 4C). Each subclusters had its own unique expression pattern. In the subclusters with high CSF1R expression, the following DEGs with significance were identified: CEBPD and IL1R2 in subcluster 1, BIRC5 and ASPM in subcluster 3, and CTSK and CKB in subcluster. On the contrary, in CSF1R subcluster with low expression, ITGB8 and PRG4 was found to be overexpressed in subcluster 2, whereas subcluster 5 overexpressed CLEC4C and LILRA4 (figure 4D). The heatmap showed the detailed patterns of DEGs in five subclusters (figure 4E). GO enrichment analysis was performed on the DEGs in five subclusters to verify their possible functions (figure 4F). According to the enrichment results, subcluster 1 was mainly related to nuclear division and organelle fission; subcluster 2 was enriched in extracellular matrix organization and responses to reactive oxygen species; subcluster 3 was enriched in chromosome segregation; subcluster 4 was associated with co-translational protein targeting to membrane; meanwhile, subcluster 5 was related to the negative regulation of apoptotic signaling pathway. In conclusion, CD163^+^CSF1R^+^ macrophages may interact with CSF1^+^ fibroblasts via the CSF1-CSF1R axis, driving the pathogenesis of PVNS.

### PVNS fibroblasts recruited the macrophages and promoted them differentiation into M2 type

To further determine the connections between fibroblasts and other cell types, cellcell communication analysis was performed (figure 5A). Compared with the other cell types, fibroblasts had the strongest cell-cell communication with macrophage. To determine the function of their connections, enrichment analysis was performed on the interacting gene pairs. Gene pairs (n=53) on the cell-cell communication axis from macrophages to fibroblasts were mainly enriched for the positive regulation of signaling pathways involving cell surface receptors, regulation of cellular component movement, regulation of cell migration and regulation of cell motility; whereas the cell-cell communication axis (n=48) from fibroblasts to macrophages were enriched in cell surface receptor signaling pathway, signal transduction, regulation of processes involving the immune system and regulation of cell migration (figure 5B, 5C).

**Figure 5.**
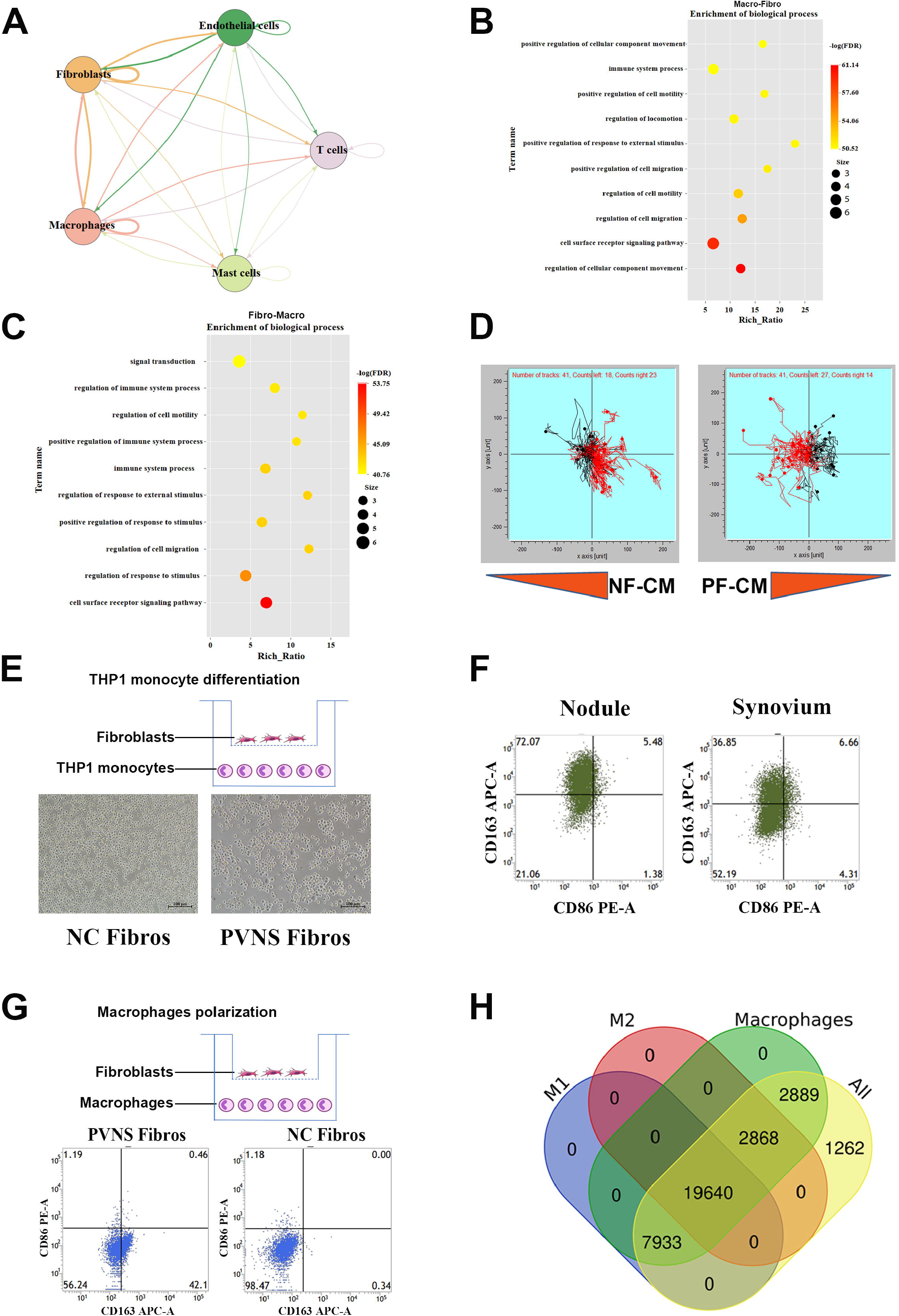
PVNS fibroblasts recruited the macrophages and promoted them differentiation into M2 type. (A) Circle plot showed the relationships among different subclusters. The arrows indicated the direction from ligands to receptors, while the thickness of the lines indicates the number of connections. (B) Results of enrichment analysis on gene pairs related to the connection between ligands of macrophages and receptors of fibroblasts in the GO: Biological Process database. (C) Results of enrichment analysis on gene pairs related to the connection between ligands of fibroblasts and receptors of macrophages in the GO: Biological Process database. (D) The result of migration assay in different fibroblasts. The trajectory of cells migrating to condition medium were marked in red. NF, negative control fibroblasts. PF, PVNS fibroblasts. CM, condition medium. (E) Schematic representation of THP1 monocytes co-cultured with fibroblasts. Scale bar, 50 μm. (F) Flow cytometry showed the different proportion of macrophages in the nodular and synovial samples of PVNS. CD163 was the marker of M2 macrophages; while CD 86 was the marker of M1 macrophages. (G) Schematic representation of macrophages co-cultured with fibroblasts. The different proportion of macrophages is reflected by the bottom plots of flow cytometry. CD163 is the marker of M2 macrophages; while CD 86 is the marker of M1 macrophages. (H) Venn plot showed the marker genes in M1 and M2 macrophages. M1, the union of macrophages expressing CD80, CD81 and CD38 genes. M2, the union of macrophages expressing CD163 gene. Macrophage, the union of CD68 positive M1 and M2 cells. All, all cells in the macrophage cluster.

Subsequently, a series of *in vitro* experiment was performed to verify the cell-cell communications between macrophages and fibroblasts, and their respective functions. In the migration experiment, monocytes were more attracted to PF-CM (conditioned medium produced by PVNS fibroblasts) than to the control medium, however, such effect was not observed in NF-CM (conditioned medium produced by normal fibroblasts) (figure 5D). Transwell co-culture assays showed that PVNS fibroblasts could promote the redifferentiation of THP1 monocyte (figure 5E). Other than that, macrophages in both the synovium and nodule of PVNS were found to be mainly M2 macrophages (figure 5F, S3A). Furthermore, when the PVNS and normal fibroblasts were co-cultured with macrophages respectively, results showed that PVNS fibroblasts induced more macrophages to differentiate into M2 type than the normal fibroblasts (figure 5G, S3B). The analysis of expression differences between M1 and M2 macrophages also showed that 7,933 genes were specifically overexpressed in M1 macrophages while 2,868 genes were overexpressed in M2 macrophages. Nevertheless, both macrophages shared 19,640 genes between them (figure 5H). These results indicated that the probable role of M2 macrophages in the occurrence of PVNS, with evidence pointing to the occurrence of nodules and poor outcome possibly due to the high proportion of M2 macrophages. However, the M1/M2 system itself is an oversimplification and may not reflect the overall complexity of macrophages.

### The cell-cell communication between fibroblasts and macrophages may lead to the development of PVNS

A series of cell communication analyses was performed between the macrophage and fibroblast subclusters to verify the function of their connection. Reannotating the UMAP plot, it was noticed that most of the fibroblasts and macrophages showed a tendency to be distributed separately, except the fibroblasts subcluster 7. The cell-cell communication analysis showed that the interactions between macrophage subcluster 1, 3 and fibroblast subcluster 3, 8 were significantly higher than the other subclusters. Meanwhile, the interactions in fibroblast subclusters between 1, 2, 3 and 8 were stronger, which have more communication connections (figure 6A). Furthermore, the connection patterns in several cell subclusters were different. For example, the connections from macrophages to fibroblasts were mainly enriched in CD74-related pathway, whereas the connections from fibroblasts to macrophages involved the FN1 pathway (figure 6B). Afterwards, CXCR and its ligands were investigated in different subclusters of cell types to determine the potential methods of connection among tumor, fibroblasts and macrophages. CXCL12 was mainly expressed in fibroblasts (fibroblasts subcluster1, 2, 3, 4 and 8), and the highest CCL5 expression was also found in fibroblasts (fibroblasts subcluster 8). Meanwhile, CXCR4 and ACKR3 were mainly expressed in macrophages (subcluster1 and 5, subcluster1 and 3, respectively) (figure 6C). These results suggested that fibroblasts and macrophages were strongly connected and may be associated with the progression of PVNS through a variety of pathways. Fibroblasts producing CXCL12 may accelerate the migration of macrophages to form tumor-like mass (figure 6D).

**Figure 6.**
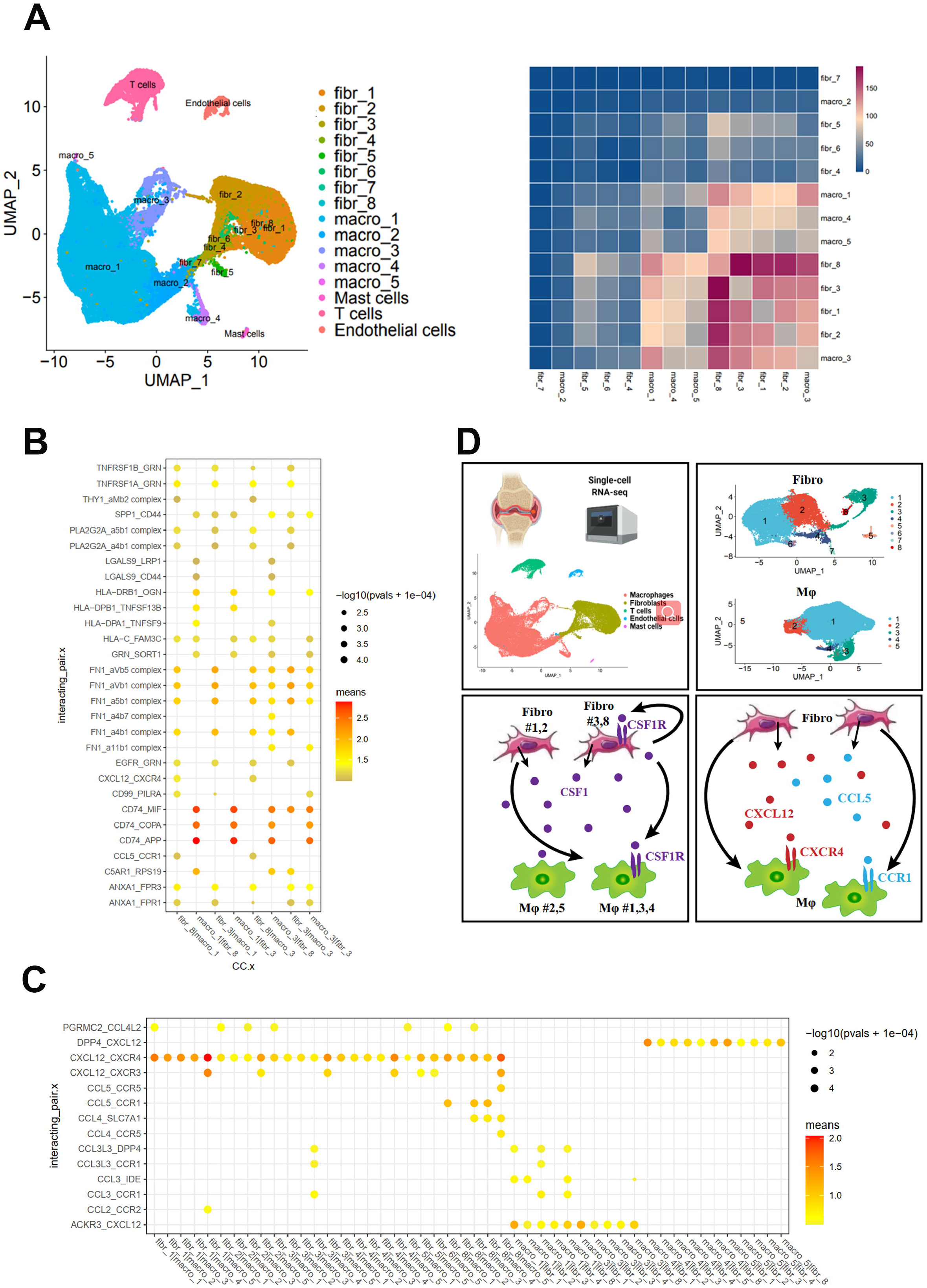
Communications among fibroblasts, macrophages and tumor cells. (A) The subcluster of fibroblasts and macrophages that were visually represented as UMAP plots. Heatmap of communications between fibroblasts and macrophages. (B) (C) The type of ligands and receptors axes in the communications between different subclusters of fibroblasts and macrophages. (D) Interaction patterns between fibroblasts and macrophages.

## Discussion

The CSF1-CSF1R axis is central in the development of PVNS, leading to the founding of several systemic therapies^13,14,22–24^. However, the exact cells that overexpress CSF1 are still not identified, nor is the specific mechanism of CSF1 overexpression elucidated. While systemic therapies of PVNS have shown promising effects under certain indications, it does not work for everyone, with some patients suffering from relapse after treatment^24^. Hence, it is crucial to figure out the complex and diverse composition of PVNS so that treatment results may be improved to benefit more patients. In this current study, scRNA-seq was performed to depict the cellular diversity of PVNS and further reveal the crosstalk between and within the components. Some specific population of fibroblasts was observed to overexpress CSF1, suggesting its vital role in the progression of PVNS. Hence, targeting the selected fibroblasts might provide a new strategy in interrupting the development of PVNS.

Although many studies have been devoted to define the cellular composition of PVNS, it remains unclear. The cellular components of PVNS include fibroblasts, macrophages (multinucleated giant cells), lymphocyte (T cells, B cells, NK cells) and monocytes, which were mainly determined by H&E staining, IHC staining, IF staining and flow cytometry^7,25–27^. Here, we comprehensively dissected the composition of localized and diffuse PVNS by scRNA-seq. From our findings, the cellular components of localized and diffuse PVNS were overall similar and consist of five clusters including macrophages, fibroblasts, T cells, endothelial cells and mast cells. Berger et al. also confirmed the similarity between both types of PVNS, highlighting the concept that localized and diffuse PVNS were two forms of one disease^26^. While the synovium and nodule of PVNS differ in morphology, they share similar composition, which suggests similar origins. Eight subclusters of fibroblasts and five subclusters of macrophages were identified, and their proposed roles in the development of PVNS were determined by the GO and KEGG analysis of each subclusters. PVNS has four key characteristics, namely pigmentation, villus, nodule and synovitis. Although the pathogenesis of these characteristics remains unclear, the functions of marker genes in each subcluster of macrophage might provide some useful clues. Further experiments are needed to explore the correlation between different subclusters and phenotypes.

The genes that encode receptor tyrosine kinases (RTKs) or their ligands are often altered in neoplastic cells, and the CSF1-CSF1R axis plays a vital role in the development of most tumors. Robert et al. depicted a tumor-landscaping effect in which the CSF1-overexpressing neoplastic cells recruit nonneoplastic cells to form a tumorous mass in PVNS. This is the basis for various PVNS systemic therapies^11^. And in 2019, FDA (the United States Food and Drug Administration) approved pexidartinib for the treatment of symptomatic PVNS. While several patients obtained satisfactory therapeutic effect, a significant number of patients experienced relapse after drug withdrawal or did not respond well at all^14,28^. Therefore, it is of importance to clarify the underlying mechanism of the CSF1-CSF1R axis in PVNS, so that other therapeutic targets may be identified. High-resolution analysis in this study confirmed that fibroblasts were the main type of cells secreting CSF1, especially fibroblast subclusters 1, 2, 3 and 8, while macrophages were the main type of cells expressing CSF1R, especially subclusters 1, 3 and 4. Moreover, compared with control group, CST7^+^CSF1^+^ fibroblasts were specific to PVNS only, suggesting their key role in the development of PVNS that was induced by the CSF1-CSF1R axis. In addition to fibroblasts, mast cells also overexpress CSF1 though fewer in number, indicating its possibly important role in the development of PVNS. Mast cells are known for their role in the inflammatory response, thus the occurrence and development of PVNS synovitis may be related to them. Further studies are needed to clarify the role of mast cells in the development of PVNS.

The role of the tumor microenvironment (TME) in tumor growth and metastasis has received more and more attention in recent years. Among the various cellular components of TME, macrophages and fibroblasts are considered as the key ones of this niche^29,30^. The role of fibroblast-macrophage interaction that centers around the CSF1-CSF1R axis has been confirmed by previous studies to be crucial in the development of many different tumors^31^. Thus, exploring the additional molecular programs that link these cells may provide novel targets for the treatment of PVNS. The cell communication analysis in this study showed that fibroblasts had the strongest interaction with macrophages. It was further confirmed that, compared to the control group, the fibroblasts of PVNS promoted a greater level of monocyte migration and differentiation. The CellPhoneDB algorithm identified the ligand-receptor interactions between CST7^+^CSF1^+^ fibroblasts and BIRC5^+^CSF1R^+^ macrophages as being the most significant. Among the ligand-receptor pairs connecting fibroblasts and macrophages, CXCL12-CXCR4 exhibits the most significant effect on promoting macrophage migration and invasion. Further studies focusing on the CXCL12-CXCR4 pair as treatment target may yield good results in tackling recurrent and refractory PVNS.

To date, macrophages are confirmed to promote cancer initiation and progression through stimulating angiogenesis, promoting metastasis, intravasation and invasion of tumor cells^32^. The polarization of macrophages is a dynamic process that can be diverted into two distinct phenotypes according to environmental stimuli and signals: the classically activated macrophage (M1 type, pro-inflammatory macrophage) and the selectively activated macrophage (M2 type, anti-inflammatory macrophage)^33^. In this current study, macrophages were confirmed to make up the most significant proportion of the PVNS tumor mass with a majority of them being the M2 type. This finding highlighted the pro-tumor role of M2 macrophages. Moreover, PVNS fibroblasts promote more macrophages toward M2 polarization than control fibroblasts. This piece of evidence suggested the vital role of fibroblasts in the progression of PVNS. However, further analysis of the scRNA-seq data demonstrated that a large number of macrophages expressed both M1 and M2 gene markers, and there were numerous macrophages that expressed neither marker. This finding suggested an oversimplification of the M1 and M2 system, thus its inability in reflecting the full complexity of macrophages. The subsets of macrophages were subdivided to further analyze the functions of each subgroup, in the hopes of clarifying the role of macrophages in PVNS. GO analysis indicated that different subsets of macrophages might pose an impact on the pigmentation, villonodule and synovitis of PVNS differently. Further investigation of the underlying mechanisms of macrophages in the four characteristics of PVNS is needed.

In conclusion, single-cell sequencing provides a comprehensive understanding of the cellular composition of PVNS, which was found to consist of macrophages, fibroblasts, T cells, endothelial cells and mast cells. Among them, CSF1-expressing fibroblasts appear to be tumor-like cells that attract macrophages to form tumor-like mass in PVNS. Furthermore, the different subsets of macrophages may play a role in pigmentation, villonodule and synovitis. These discoveries provide hopeful insights for the treatment of PVNS by targeting fibroblasts and their related ligand receptor pairs.

## Acknowledgements

We would like to thank Phei Er Saw for proofreading and language improving this article. This study was financially supported by grants from the National Natural Science Foundation of China (81972067), the National Natural Science Foundation of China (81802127) and the Health Welfare Fund Project of Futian District (FTWS2020078).

---

**Figure S1. Clinical characteristics of the three PVNS patients and the cellular composition of six samples**

(A) The quality control of the scRNA-seq samples.

(B) Demographic and clinical characteristics of patient samples used in scRNA-seq.

(C) The cellular composition of the six scRNA-seq samples.

**Figure S2. scRNA-seq defines the cellular ecosystem of PVNS**

(A) Identical UMAP demonstrating the marker gene expression levels of different cell types.

(B) Cellular composition of the six scRNA-seq samples.

(C) Macrophage-cell-type fractions relative to the total macrophage cell count per sample.

(D) Fibroblast-cell-type fractions relative to the total fibroblast cell count per sample.

(E) Proportion of fibroblasts from PVNS or control sample within each cluster.

**Figure S3. Flow cytometry gating strategy**

(A) Flow cytometry gating strategy for macrophage populations in PVNS sample.

(B) Flow cytometry gating strategy for macrophage populations from PBMC.

