## Supplementary figures and images for "Single-cell RNA-seq analysis reveals aberrant CSF1 expression in disease-causing synovial fibroblasts of pigmented villonodular synovitis"

### figure S1

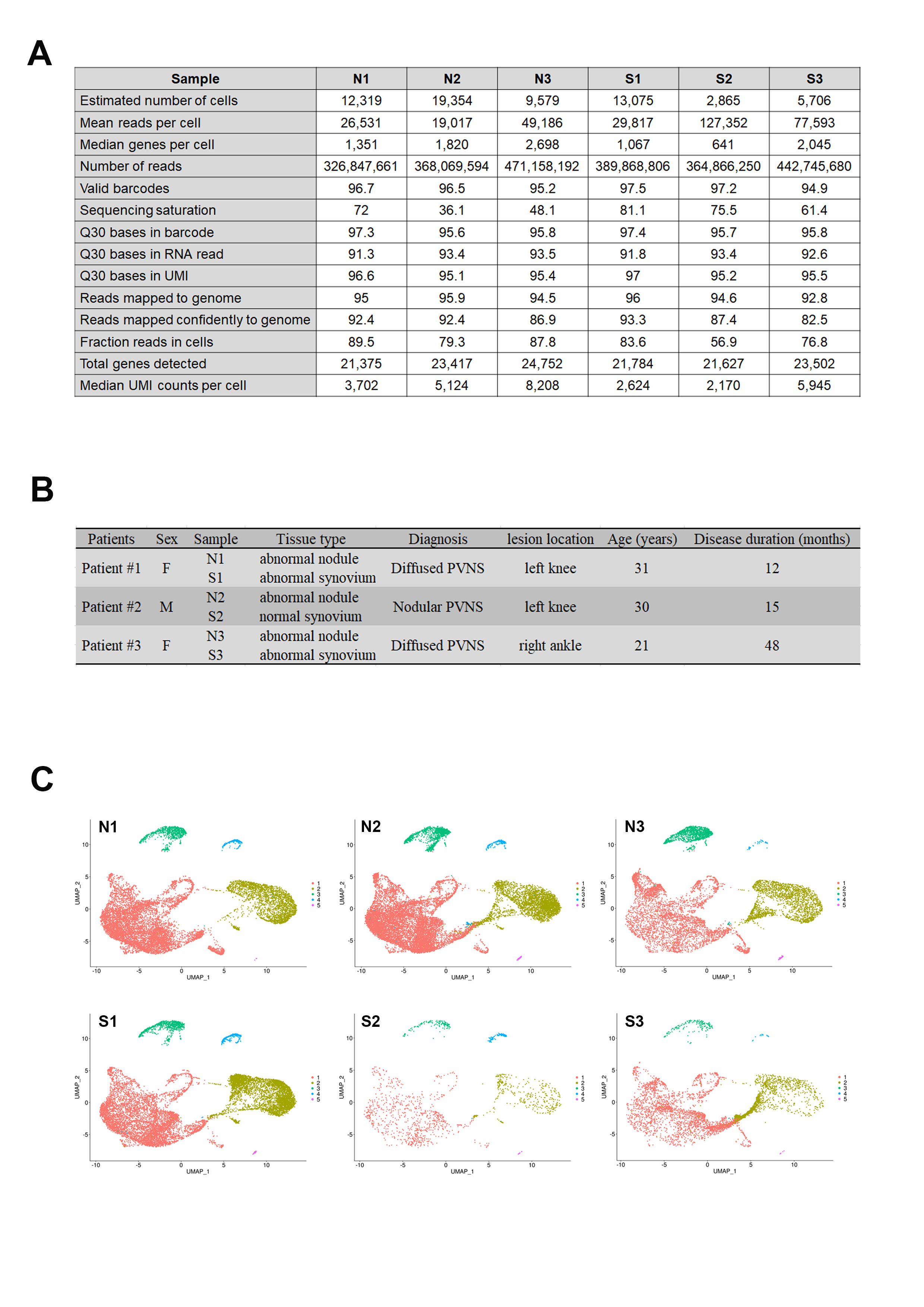

### figure S2

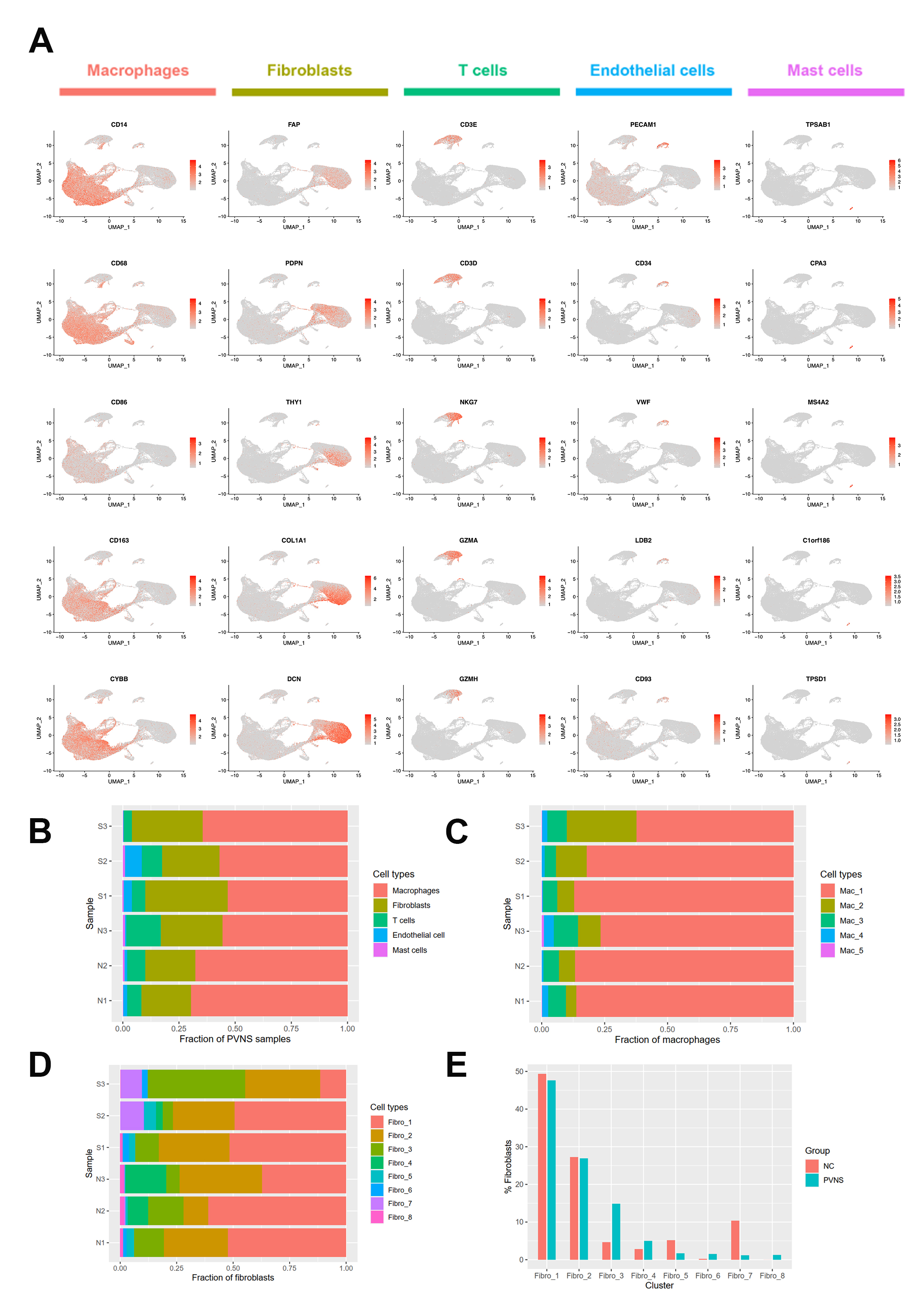

### figure S3

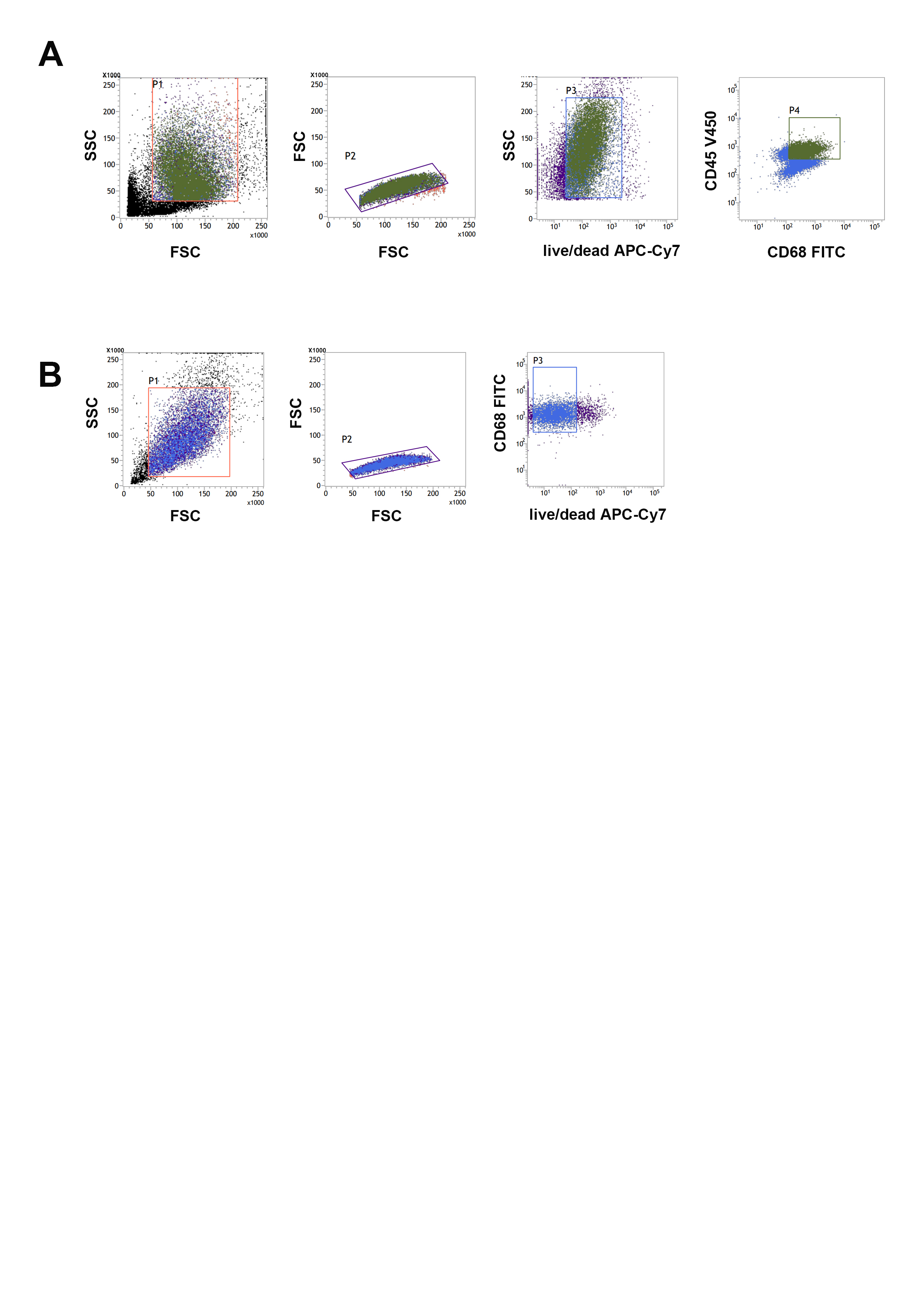
